# Enterobactin carries iron into *C. elegans* and mammalian intestinal cells by a mechanism independent of divalent metal transporter DMT1

**DOI:** 10.1101/2024.12.20.629725

**Authors:** Aileen K. Sewell, Mingxue Cui, Mengnan Zhu, Miranda R. Host, Min Han

**Affiliations:** Department of MCDB, University of Colorado Boulder For correspondence

**Keywords:** <u>Keywords</u>: smf-3, Ent, Enterochelin, FeEnt, anemia, iron deficiency, iron transport, iron metabolism, siderophore, hif-1, hypoxia-inducible factor, ferritin, ferroportin, ftn-1, ftn-2, fpn-1.2

## Abstract

The diverse microbiota of the intestine is expected to benefit the host, yet the beneficial metabolites derived from the microbiota are still poorly understood. Enterobactin (Ent) is a well-known secreted iron-scavenging siderophore made by bacteria to fetch iron from the host or environment. Little was known about a positive role of Ent until a recent discovery in the nematode *C. elegans* indicated a beneficial role of Ent in promoting mitochondrial iron level in the animal intestine. To solidify this new paradigm, we further tested this role in *C. elegans* and multiple mammalian cell models and its relationship with the primary iron transporter DMT1/SMF-3 and several other iron-related genes. Here we show that ferric enterobactin (FeEnt) supplementation promotes whole organism development in *C. elegans*, increases iron uptake in caco-2 human intestinal epithelial cells, and supports iron-dependent differentiation of murine erythroid progenitor cells, indicating that the FeEnt complex can effectively enter these cells and be bioavailable. Our data in multiple models demonstrate that FeEnt-mediated iron transport is independent of all tested iron transporters. In addition, FeEnt supplementation robustly suppresses the developmental defects of a *hif-1* mutant under low iron condition, suggesting the critical role in iron homeostasis for this well-known hypoxia regulator. These results suggest that FeEnt can effectively enter animal cells and their mitochondria through a previously unknown mechanism that may be leveraged as a therapeutic ferric iron carrier for the treatment of DMT1- or HIF-1-related iron deficiency and anemia.

## Introduction

Enterobactin (Ent), a siderophore with the strongest known affinity for ferric iron (1), is naturally produced by certain bacteria such as *E. coli*. This non-ribosomal peptide is synthesized by a multistep process within the bacteria, then excreted into the environment to scavenge iron under low-iron conditions to benefit bacterial survival (2, 3). After binding to Fe^3+^, the ferric enterobactin (FeEnt) complex is transported back into the bacteria where iron is released and utilized.

In the FeEnt complex, Fe^3+^ is tightly bound by six hydroxyls from the three catecholate groups of Ent, making it a highly stable and negatively charged structure (4–6). FeEnt enters bacterial cells through a specialized receptor system, needed at least in part to overcome the bacterial cell wall (7). However, whether and how Ent or FeEnt enter animal cells is not clear. Citing the molecular weight, negative charge, and other properties of the complex, some siderophore experts considered it highly unlikely that Ent could passively enter animal cells and thus be used to transport iron into animals (7–11).

Using innovative assays and an unbiased genetic screen, our lab previously discovered an unexpected role of Ent to support growth and the labile iron pool in *C. elegans* that is independent of the bacterial usage of Ent (12). Ent dietary supplementation was shown to increase the iron level in *C. elegans’* mitochondria and did so in a manner that was dependent on its binding to the ATP synthase α-subunit (ATPSα). Additional experiments also indicated that Ent can transfer iron into the mitochondria of mammalian cells and such activity is also likely dependent on the interaction with ATPSα (12). However, how FeEnt enters animal cells and their mitochondria is still an outstanding question.

Divalent Metal Transporter 1 (DMT1) is located on the apical membrane of enterocytes where it plays a major role in ferrous iron (Fe^2+^) uptake from the diet into the cell (13–15). While DMT1 only mediates the transport of Fe^2+^ iron, dietary Fe^3+^ ions may be reduced to Fe^2+^ by duodenal cytochrome b (DCYTB) at the outer cell membrane. In addition, DMT1 is also known to transport iron from the endosome/lysosome to the cytosol where it is available for mitochondrial iron uptake(13, 15), and has also been found on the mitochondrial outer membrane (16, 17). DMT1 is a critical player in iron homeostasis. Whether DMT1 is involved in Ent-mediated cellular and mitochondrial iron uptake in animals is a significant question to address.

In this study, we utilized an iron-deficient model in *C. elegans* to demonstrate the effectiveness of Ent as a vehicle for iron uptake into animal cells. Through this pursuit we discovered that the FeEnt benefit does not rely on DMT1/SMF-3 transport activity, indicating an alternative mechanism for FeEnt uptake. We found that this DMT1-independent mechanism is also conserved in mammals.

## Results

### Ent promotes iron uptake in *smf-3*(-) *C. elegans*

Our previous study demonstrated the surprising benefit of Ent to the physiology of the nematode *C. elegans* by increasing iron levels and promoting growth (12). Our further studies have focused on understanding the mechanism by which Ent brings iron into animal cells. One possible model is that an unknown transporter or protein interacts with and facilitates the import of FeEnt. Although the Ent-ATPSα interaction was shown to be critical for the beneficial role, such an interaction was shown to be dispensable for FeEnt to be transported into mitochondria (12). Another important candidate protein is DMT1. Ubiquitously expressed throughout the body, DMT1 is prominently expressed and localized to the apical membrane of enterocytes where it plays a major role in iron absorption from the intestine (18). We thus tested if the DMT1 function is required for the Ent benefit in *C. elegans*. The nematode *C. elegans* has three genes that are homologous to DMT1, *smf-1/2/3*.

These three genes have unique expression patterns, with *smf-1* and *smf-3* expressed in different regions of the intestine, while *smf-2* is expressed in the pharynx, gonad, and neurons (19).

Measured by different methods, both *smf-3*(-) and *smf-2*(-) have been reported to have reduced iron levels (19, 20). Although the three *smf* mutants individually lack strong phenotypes under standard laboratory conditions (19), *smf-3*(-) was recently shown to exhibit developmental delay under low-iron growth conditions (21), suggesting that SMF-3 is the key iron transporter in the worm intestine and its mutant defect in iron homeostasis cannot be sufficiently compensated for by the functions of the other two SMF proteins. We utilized this iron-poor *smf-3*(-) developmental delay phenotype as an assay to test if the Ent benefit to animal growth required SMF-3/DMT1. Briefly, *smf-3*(-) mutants were hatched on OP50 NGM agar plates containing the iron chelator 2,2’-bipyridyl (BP) (“OP50/BP assay” hereafter) and scored for developmental stage (Fig. 1A). Although *smf-3*(-) worms show no growth delay on standard lab conditions (OP50/NGM), the *smf-3*(-) mutant worms cultured under this iron-poor condition displayed a significant developmental delay. We found that this growth delay was significantly suppressed by the addition of Ent, supporting growth to fertile adulthood (Fig. 1B). These results suggest that Ent benefits worm growth under iron-poor conditions, and does so by an SMF-3- independent mechanism.

**Figure 1.**
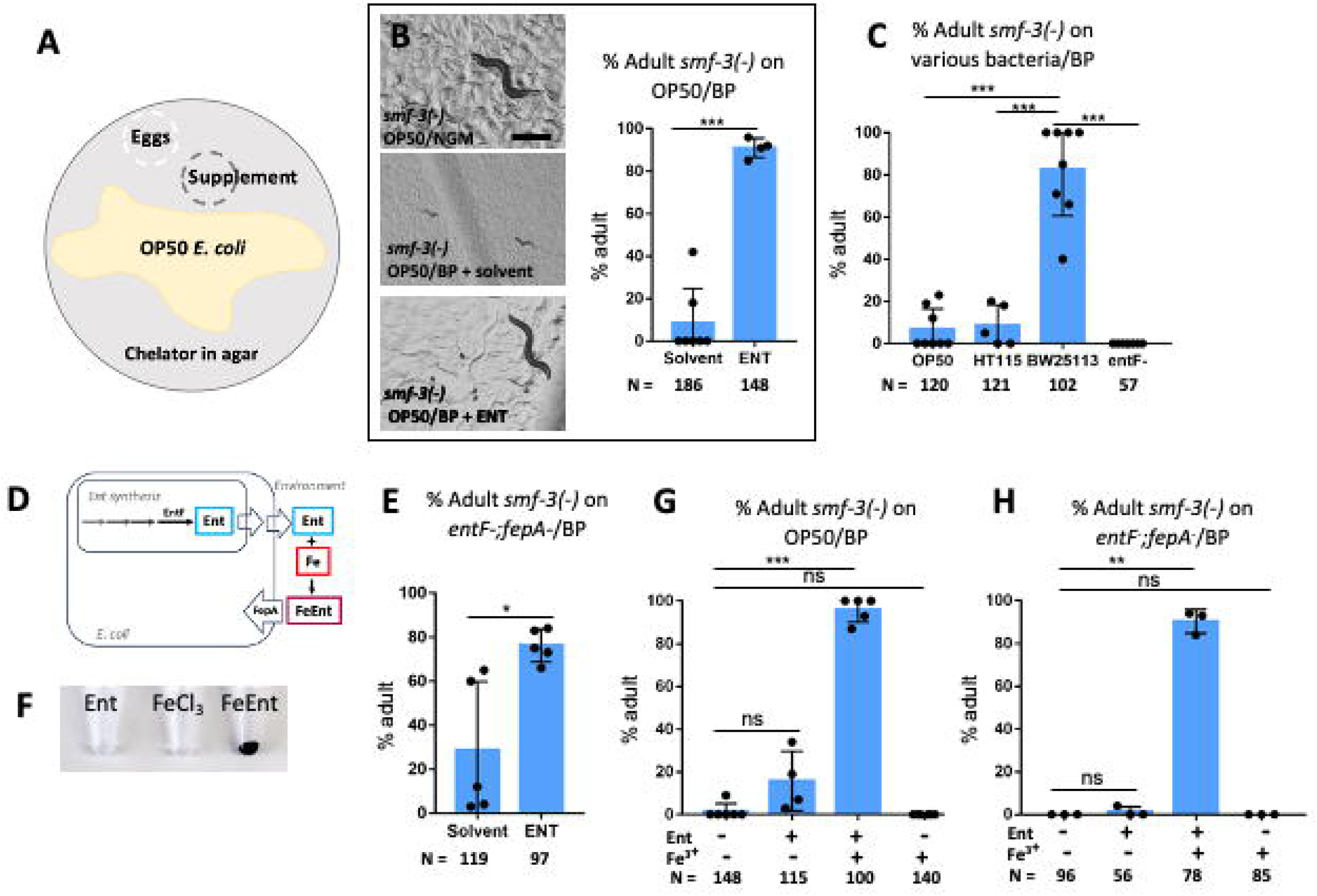
**Ent benefits *smf-3(-)* worm growth under iron-poor condition**. (A) Diagram of OP50/BP worm assay plate. Agar contains the chelator 2,2’-bipyridyl (BP). (B) Representative images and bar graph of *smf-3(-)* worm growth to adult when supplemented with or without Ent. Ent supplementation significantly rescued worm development to gravid adults, similar to the standard (NGM) growth condition. Solvent is DMSO. NGM: nematode growth media, no chelator. Scale bar equals 500 microns. The difference between treatment groups on day five was found to be statistically significant. (C) Bar graph of *smf-3(-)* worm growth to adult when fed different bacteria. Strains OP50 and HT115 do not support *smf-3(-)* growth on the BP chelator condition. BW25113 does support growth, but a single mutation in entF (Ent synthesis gene) eliminates the effect. The growth effect seen with BW25113 was found to be statistically different from OP50, HT115, and entF- tests on day five. (D) Simplified diagram of Ent synthesis and uptake pathways in *E. coli*. (E) Bar graph of *smf-3(-)* worm growth to adult when fed *entF-;fepA-* mutant bacteria that could neither produce nor take up Ent, supplemented with or without Ent. *smf-3(-)* growth was significantly increased with Ent supplementation compared to solvent control on day five. (F) Image of strong color change that quickly occurs when equimolar solutions of FeCl_3_ and Ent are combined to form FeEnt. (G) Bar graph of *smf-3(-)* worm growth to adult when supplemented with Ent, FeCl_3_, or FeEnt. FeEnt significantly rescued worm development (compared to control) and did so more rapidly than Ent alone, on day four. (H) Bar graph of *smf-3(-)* worm growth to adult when fed *entF-;fepA-* mutant bacteria and supplemented with Ent, FeCl_3_, or FeEnt. FeEnt significantly rescued worm development (compared to control) when fed the *entF-;fepA- E. coli* strain that could neither produce nor take up Ent, on day four. N = total number of worms scored. Error bars are standard deviation. Individual data points represent independent biological replicates. ∗*p* < 0.05, ∗∗*p* < 0.01, ∗∗∗*p* < 0.001, ns: not significant; unpaired two-tailed Student’s *t* test.

### Ent supplied by *E. coli* can also benefit the growth of *smf-3*(-) *C. elegans*

In the above tests, the presence of the BP chelator created a relatively low iron condition that might be expected to induce the *E. coli* to produce more Ent. However, the data in Fig. 1B suggested that the OP50 *E. coli* strain did not produce Ent to support animal growth. This theory is consistent with a previous report that the OP50 *E. coli* strain lacks the biosynthetic enzymes required to produce Ent (22). To determine if Ent supplied by *E. coli* could also benefit *C. elegans* lacking the SMF-3/DMT1 transporter, we first tested if other *E. coli* strains could support *smf-3*(-) growth on the BP condition. We found that while *smf-3*(-) mutants grew poorly on standard lab strains like OP50 and HT115 when combined with the BP chelator, there was no growth defect when fed with a K12 *E. coli* strain (BW25113) (Fig. 1C), suggesting that K12 BW25113 produces Ent that benefits worm growth. We then took advantage of the Keio collection, an *E. coli* single mutant library based on the K12 BW25113 strain. To determine if the K12 BW25113 growth benefit was specifically due to Ent, we tested *smf-3*(-)/BP growth on the *entF-* single mutant that cannot synthesize Ent, and found *entF-* did not support *smf-3*(-) growth (Fig. 1C-D). EntF is an essential gene for the biosynthesis of Ent in *E. coli* [Fig. 1D; (2, 3)]. These results clearly indicate that Ent synthesized by the K12 *E. coli* strain promotes the growth of *smf-3*(-) animals.

### Ent benefit to the host is independent of Ent-related metabolism in *E. coli*

We next tested if the Ent supplementation was benefiting the worm directly, or benefiting the worm indirectly through an *E. coli*-dependent mechanism. Although the OP50 *E. coli* strain lacks the Ent biosynthesis enzymes (22), it may still benefit from the Ent supplement on the plate and take up ferric Ent by FepA, the ferric Ent transporter (Fig. 1D). If so, OP50 *E. coli* might utilize the Ent added to the culture plate to fetch iron and improve its own physiology, which may present an indirect benefit to *C. elegans* through the diet. To address such a possibility, we repeated the test using an *E. coli* double mutant that can neither synthesize nor take up Ent (BW25113 *entF-; fepA-)* (Fig. 1D). We found that Ent supplementation was still effective to significantly suppress the growth defect of *smf-3*(-) mutant worms fed the *entF-; fepA*- double mutant *E. coli* in the presence of BP (Fig. 1E), indicating that the benefit of Ent supplementation under this condition is independent of Ent-related activities in bacteria. Therefore, the Ent growth benefit shown in Fig. 1B is likely direct to *C. elegans* by an *E. coli*-independent mechanism.

### Ferric Ent is more effective and beneficial to *C. elegans* than Ent alone

Our previous work presented a model where Ent aids the host by facilitating the uptake of Fe^+3^ from the environment (12). Based on this model, we asked if supplementation with FeEnt (Ent that has been incubated with FeCl_3_ to form the Ferric-Ent complex) would also support *smf- 3*(-) mutant growth by the OP50/BP assay. Mixing equimolar solutions of Ent and FeCl_3_ to produce FeEnt caused a strong color change indicating binding (Fig. 1F). By the same growth assay, we found that FeEnt also benefited *smf-3*(-) growth on both OP50/BP (Fig. 1G) and *entF-; fepA-*/BP (Fig. 1H) growth conditions. We found that FeEnt significantly suppressed the growth defect of the *smf-3*(-) mutants (compared to control) and did so 24hrs earlier than the effect seen with Ent supplementation alone (Fig. 1G-H). Figure 1G-H represent the day four growth score, when the majority of Ent-treated worms under these assay conditions do not reach adulthood until day five (Fig. 1B). These results support the model that Ent benefits *C. elegans* by binding and transporting ferric iron into animal cells, and Fe-Ent benefits the iron-deficient host at a lower dose than FeCl_3_ supplementation alone.

### FeEnt benefit to host is independent of key iron transport genes

The robust developmental phenotype demonstrated by the *smf-3*(-) OP50*/*BP assay, and the strong rescue by FeEnt, suggest that SMF-3 is the primary iron transporter in *C. elegans* and that FeEnt is beneficial to the host by an SMF-3-independent mechanism. We took this analysis further to address the potential role of the other SMF proteins in this beneficial mechanism. First, we created the *smf-1*(-) *smf-3*(-) double mutant, targeting the two DMT1 homologs expressed in the *C. elegans* intestine. We found that the double mutant is viable and looks superficially wildtype under standard lab conditions. We then tested the *smf-1*(-) *smf-3*(-) double mutant by the OP50/BP assay and found that the double mutant showed the same growth delay as the *smf- 3*(-) single mutant, and the double mutant growth was similarly and significantly rescued by supplementation with FeEnt, and not FeCl_3_ alone (Fig. 2A-B). Second, we knocked down all three *smf* genes. Because an *smf-1/2/3* triple mutant cannot be created by traditional cross methods due to the slight genetic distance between *smf-1* and *smf-2*, we created the *smf-2*(-)*; smf- 3*(-) double mutant and then used *smf-1(RNAi)* by feeding [SMF-1 is prominently expressed in the intestine, making it a suitable RNAi target (19)]. Similar to the *smf-3*(-) mutant, the *smf-2*(-)*; smf-3*(-) double mutant was viable and displayed normal growth under standard lab conditions, but had the same delayed growth phenotype on the OP50/BP condition. The *smf-2*(-)*; smf-3*(-) double mutant was grown from hatching on *smf-1(RNAi)* or empty vector control, then the resulting adults were bleached and eggs were placed on OP50/BP plates, with or without supplements (Fig. 2C). Developmental stage was scored five days later (Fig. 2D-E). We found that *smf-2*(-)*; smf-3*(-)*; smf-1(RNAi)* mutants had a strong developmental delay by the OP50/BP assay that was significantly rescued to fertile adults by FeEnt supplementation, but not by FeCl_3_ supplementation alone (Fig. 2D). These data further support the claim that FeEnt benefits *C. elegans* by an SMF-1/2/3-independent mechanism.

**Figure 2.**
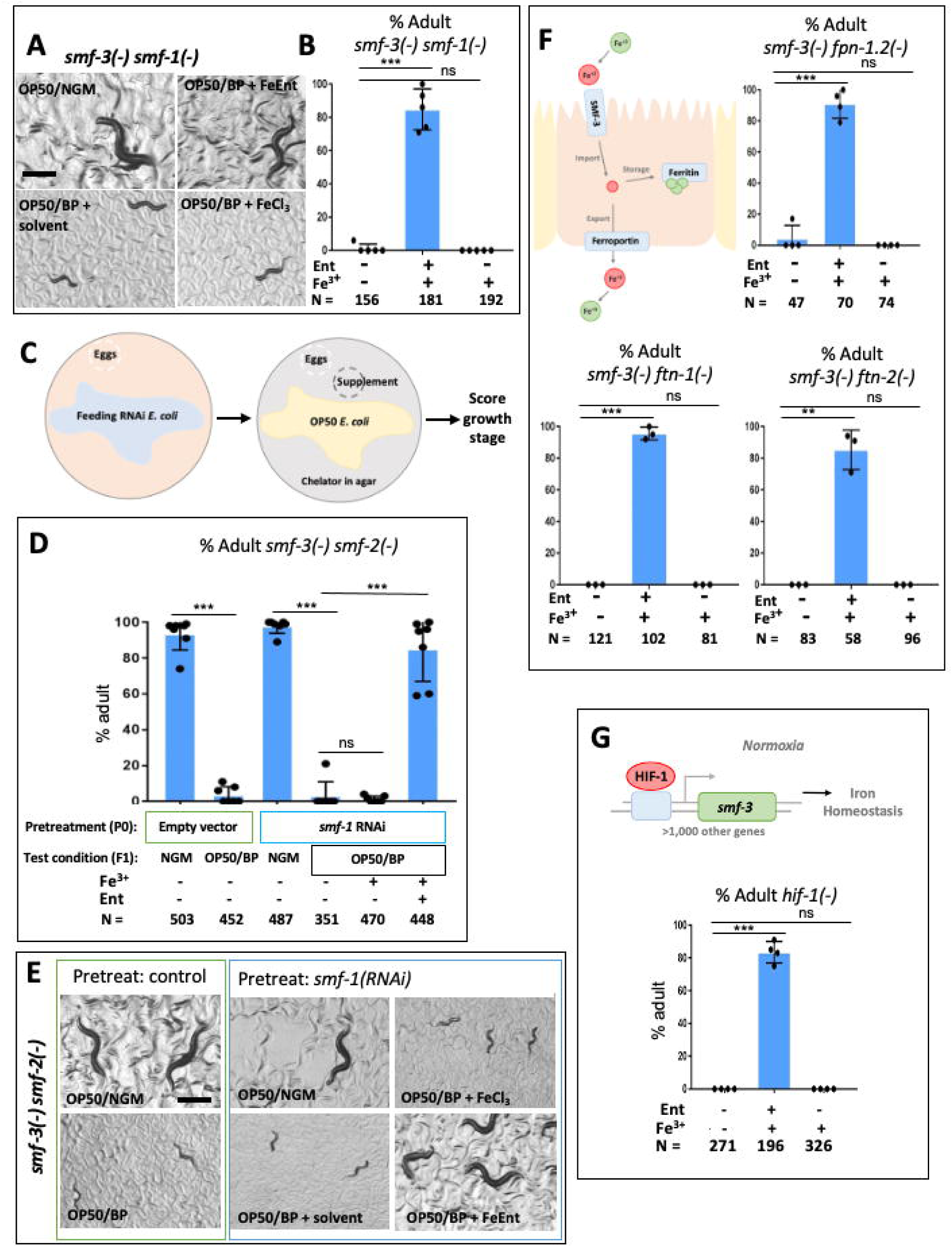
FeEnt benefit to host is independent of key iron transport and metabolism genes. (A and B) Representative images and bar graph of *smf-3(-) smf-1(-)* double mutant worm growth to adult when treated with or without supplements. FeEnt supplementation significantly rescued worm development to gravid adults, similar to the standard (NGM) growth condition. Solvent is DMSO. NGM: nematode growth media, no chelator. The difference between treatment groups on day five was found to be statistically significant. (C) Diagram of feeding RNAi assay with OP50/BP test. *smf-3(-) smf-2(-)* double mutant worms were treated on RNAi by feeding for one generation, then gravid adults were bleached to the OP50/BP assay plate, with or without supplements, and growth stage was scored five days later. (D) FeEnt supplementation significantly rescued *smf-1/2/3(-)* worm development to gravid adults, but not FeCl_3_ alone. (E) Representative images of worm growth after indicated treatment. (F) Simplified diagram of iron import, storage, and export in an intestinal cell. Bar graphs showing FeEnt supplementation significantly rescued worm development of *smf-3(-)*-containing double mutant strains to gravid adult stage, but not FeCl_3_ alone. Ferritin homologs: *ftn-1, ftn-2*. Ferroportin homolog: *fpn-1.2*. Solvent is DMSO. (G) Simplified diagram of HIF-1 regulation of gene expression under normoxia. Bar graph shows *hif-1(-)* mutants could not grow to adult on the OP50/BP assay plate, but growth was significantly rescued by supplementation with FeEnt. N = total number of worms scored. Error bars are standard deviation. Scale bar equals 500 microns. Individual data points represent independent biological replicates. ∗*p* < 0.05, ∗∗*p* < 0.01, ∗∗∗*p* < 0.001, ns: not significant; unpaired two-tailed Student’s *t* test.

We next tested the potential role for genes involved in iron storage (ferritin) and export (ferroportin) in this beneficial mechanism (Fig. 2F). The iron storage protein ferritin has two homologs in *C. elegans*. *ftn-1* is expressed in the intestine and neurons, and *ftn-*2 is expressed in all major tissues. *ftn-1(ok3625)* mutants are superficially wildtype, while *ftn-2(ok404)* mutants have been reported to have decreased iron levels. The iron export protein ferroportin has two predicted homologs in *C. elegans*, *fpn-1.1* and *fpn-1.2*, but these individual mutants have no reported phenotypes. Surprisingly, FTN-1, FTN-2, and FPN-1.2 proteins were not required for the FeEnt mechanism. The combination of each of these mutations with *smf-3*(-) was rescued by FeEnt in the OP50/BP assay, indicating that these factors individually are not required for the FeEnt benefit (Fig. 2F). These results may be due to genetic redundancy for these functions.

### FeEnt supplementation robustly suppressed developmental defect of *hif-1*(-)

*C. elegans* encode one protein, HIF-1, that is the worm homolog of the well-studied hypoxia-inducible transcription factor HIF-1α and HIF-2α in vertebrates (23). HIF-1 is primarily known for regulating gene expression for adaptation to hypoxia, but has also been surprisingly shown to regulate the transcription of 1,075 genes under normoxic conditions (24), including *smf-3* (20). Because *hif-1(ia4)* mutants are superficially healthy under standard lab and oxygen conditions, but have been shown to cause developmental delay under iron-poor conditions [similar to *smf-3*(-)](21), we tested if FeEnt could rescue their growth. We found that the *hif- 1(ia4)* mutant growth delay observed by the OP50/BP assay was significantly suppressed by the addition of FeEnt, supporting growth to fertile adulthood (Fig. 2G). This positive, beneficial result shows that FeEnt is sufficient to rescue *hif-1*(-), and may suggest that FeEnt benefits the host by a mechanism that is independent of HIF-1 regulation and/or any of the 1,075 HIF-1- regulated genes under normoxia. These data also support the critical role of HIF-1 in regulating iron homeostasis.

### Ent facilitates DMT1-independent iron transport in cultured human intestinal epithelial cells

We then tested if Ent can also carry iron into mammalian intestinal epithelial cells, and explored if this activity is dependent on the DMT1 transporter. We chose to test Caco-2 cells, a human intestinal epithelial cell line derived from human colorectal adenocarcinoma, that has been widely used as an *in vitro* model for intestinal absorption of nutrients and drugs, including iron (25). This cell line was shown to have a high level of DMT1 expression (26). In our experiments, we confirmed that ferrous iron uptake was significantly higher than ferric iron in Caco-2 cells (Fig. 3A). We then used Ebselen, a potent inhibitor of DMT1 that was originally identified in a high-throughput screen (27), to reduce DMT1 activity in Caco-2 cells. In our tests, Ebselen treatment (20uM) could significantly reduce ferrous iron uptake but had no impact on ferric iron uptake (Fig. 3B-C).

**Figure 3.**
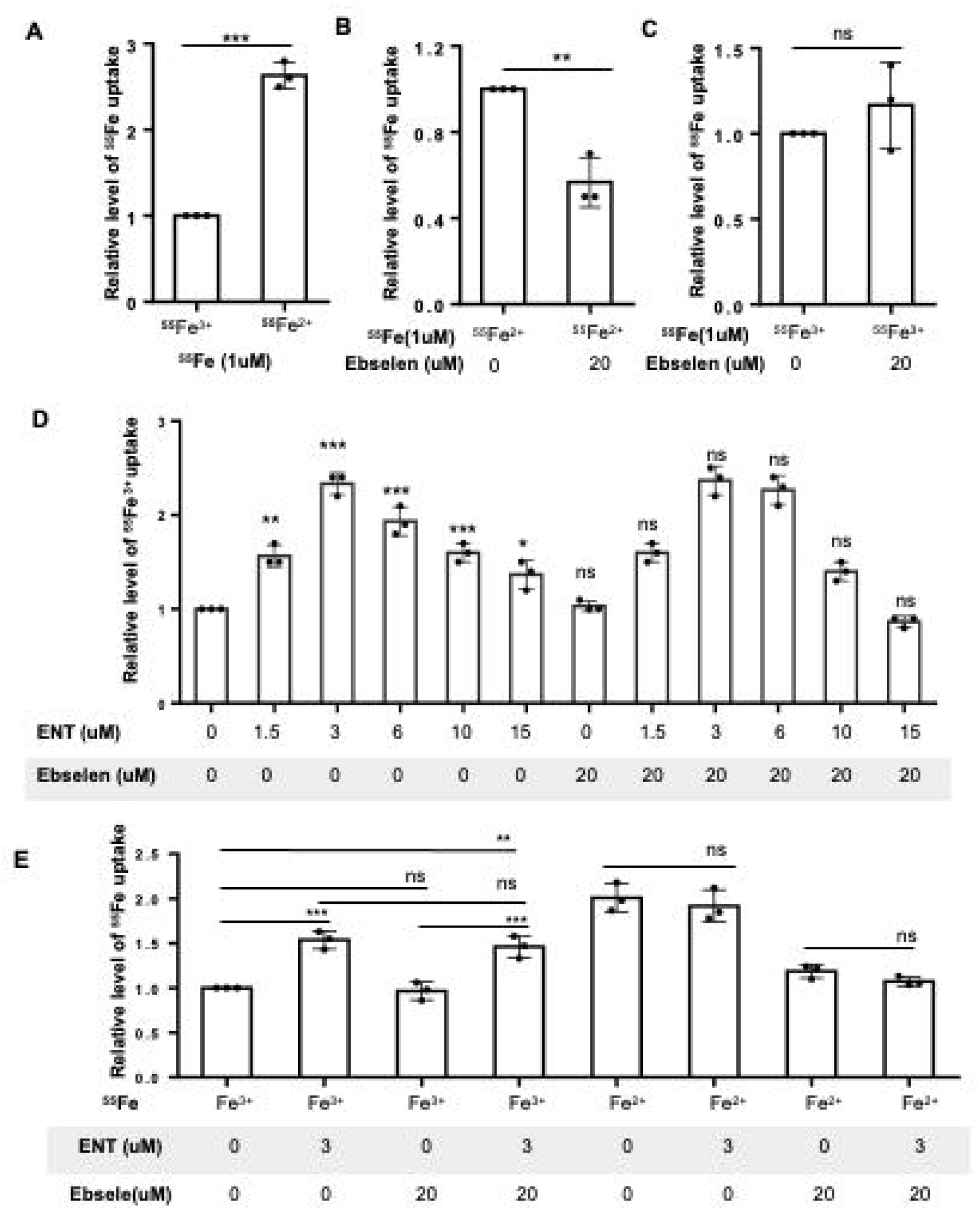
**Inhibiting DMT1 does not affect Ent-mediated iron uptake in Caco-2 cells**. Iron absorption into Caco-2 cells was evaluated by measuring the level of ^55^Fe in cells. (A) Comparison of ferric and ferrous iron uptake in Caco-2 cells. The radioactivity of the sample treated with ^55^Fe^3+^ was used as control and was set as 1. The relative level of ^55^Fe^2+^ uptake was calculated by dividing the radioactive value of cells treated with ^55^Fe^2+^ by the control radioactive value. (B and C) The effect of Ebselen (20 uM) treatment on ferrous iron uptake (B) or ferric iron uptake (C) was analyzed by comparing the radioactivity after cells treated with or without Ebselen. The radioactivity of the sample treated without Ebselen was used as control and was set as 1. The relative level of ^55^Fe uptake was calculated by dividing the radioactive value of cells treated with Ebselen by the control radioactive value. (D) The effect of Ebselen on the Ent- regulated Fe^3+^ uptake was analyzed by culturing cells with various Ent concentrations and treated with or without Ebselen. (E) The effect of Ent on Fe^3+^ and Fe^2+^ uptake without or without Ebselen treatment was analyzed. The radioactivity of the sample treated with neither Ent nor Ebselen was used as control and was set as 1. The relative level of ^55^Fe uptake was calculated by dividing the radioactive value of cells treated with either Ent or Ebselen, or both, by the control radioactive value. Error bars are standard deviation. Individual data points represent independent biological replicates. ∗*p* < 0.05, ∗∗*p* < 0.01, ∗∗∗*p* < 0.001, ns: not significant; unpaired two-tailed Student’s *t* test.

We then supplied Ent to Caco-2 cells and found that Ent exhibits a non-monotonic, biphasic effect on ferric iron uptake, suggesting an optimal concentration for maximum impact (Fig. 3D). Specifically, in the absence of Ebselen, Ent promoted the highest ferric iron uptake at 3 μM, but such an ability was compromised at either lower or higher concentrations. The cause of this effect remains undetermined. Importantly, DMT1 inhibition did not alter the ability of Ent to facilitate ferric iron transport into Caco-2 cells (Fig. 3D). We then further compared the impact of adding Ent with or without Ebselen on the uptake of ferrous iron with that of ferric iron. We found that addition of 3 μM Ent had no significant impact on the cellular ferrous iron level, which was prominently different from the impact on ferric iron uptake (Fig. 3E).

Moreover, Ebselen treatment significantly reduced ferrous iron uptake in cells with or without Ent addition (Fig. 3E). These data support the idea that Ent specifically facilitates ferric iron transport in mammalian epithelial cells and does so through a DMT1-independent mechanism.

### Ent promotes DMT1-independent iron transport in human HEK293F **DMT1 2/-IRE** cells

In addition to using Caco-2 cells and the DMT1 inhibitor Ebselen, we also tested a human HEK293F cell line that expresses a DMT1 isoform (DMT1 2/-IRE) under the control of tetracycline (including doxycycline) (28). Since HEK293F cells, derived from embryonic kidney cells, have a very low level of DMT1 expression, the inclusion of the DMT1 2/-IRE transgene permits conditional control of the DMT1 expression level. Like in Caco-2 cells, we found a similar dosage-dependent effect of Ent supplementation on ferric iron uptake in the uninduced HEK293 DMT1 2/-IRE cells (Fig. 4A). The addition of 1.5 uM Ent displayed the highest effect on ferric iron uptake, which is slightly lower than that found in Caco-2 (Fig. 3D). Upon induction of the expression of the DMT1 2/-IRE protein with the addition of doxycycline (Fig. 4B), ferrous iron uptake was significantly increased (Fig. 4C). In contrast, the doxycycline induction had no significant impact on ferric iron uptake with or without Ent treatment (Fig. 4C). This data again support that Ent mediates a DMT1-independent iron transport mechanism in mammalian cells. In addition, these data also suggest that such role of Ent is not limited to enterocytes where dietary iron is normally absorbed.

**Figure 4.**
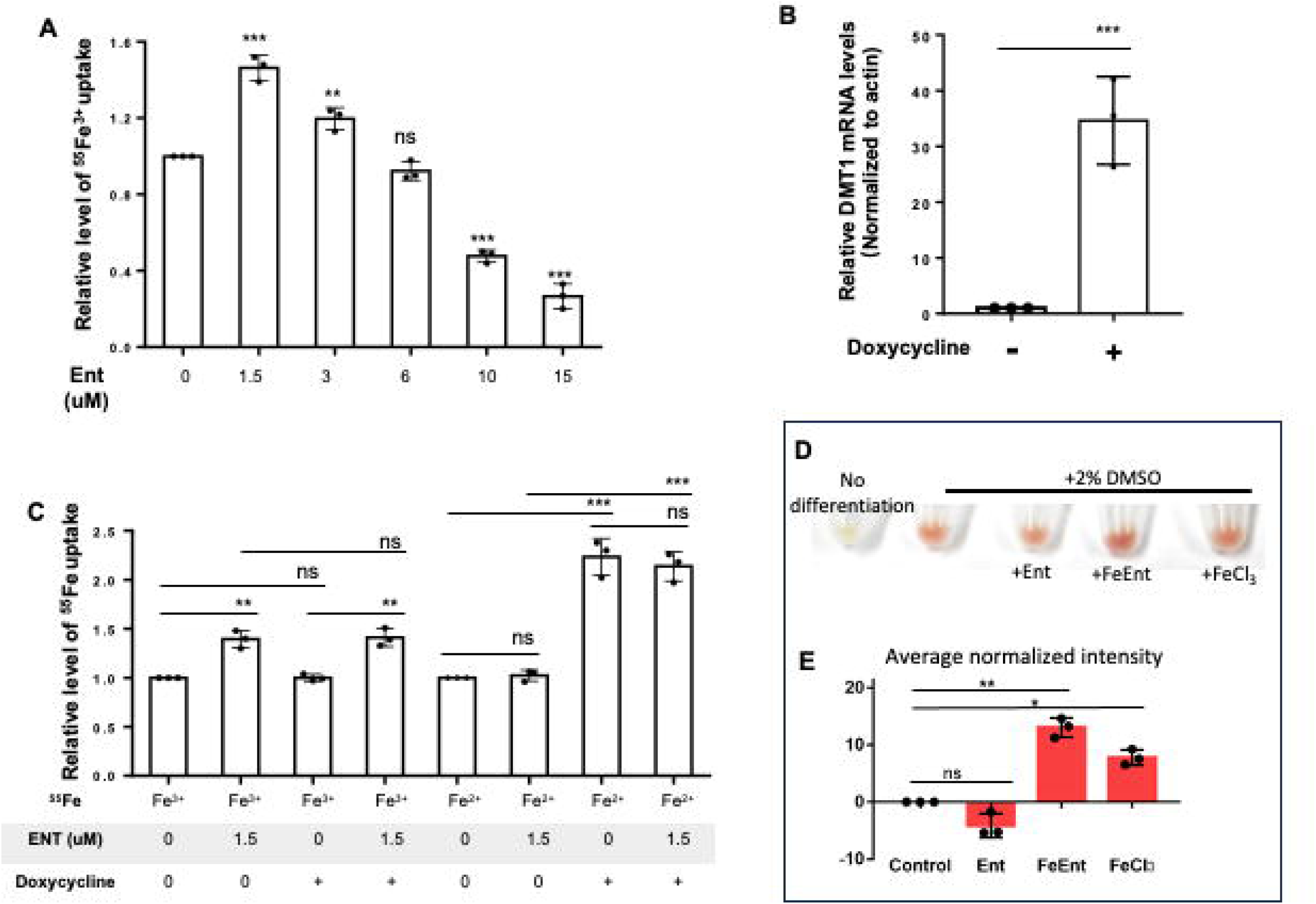
Ent promotes ferric iron uptake in human HEK293F_2/-IRE-DMT1 cells and murine erythroid progenitor cells. (A) Effect of Ent on Fe^3+^ uptake was measured in HEK293F_2/-IRE-DMT1 cells. Cells were treated with various concentrations of Ent. The radioactivity of the sample treated without Ent was used as control and was set as 1. The relative level of ^55^Fe^3+^ uptake was calculated by dividing the radioactive value of each experimental group by the control radioactive value. (B) Mouse-specific DMT1 mRNA expression levels in HEK293F_2/-IRE-DMT1 cells with (+) or without (-) exposure to doxycycline (50nM) were analyzed by qRT-PCR. Cells were grown to ∼60% confluence in 6-well plates. Either doxycycline (50 nM) or water (as control) was added, and cell cultures were grown for 24 hours. Individual data points represent independent biological replicates. (C) Role of DMT1 expression induced by doxycycline in HEK293_2/-IRE-DMT1 cells on Ent-regulated Fe^3+^ uptake. Triplicates were examined for each experimental condition. (D and E) Representative images and bar graph showing FeEnt addition promoted differentiation of murine erythroid progenitor (MEL) cells. The differentiation of MEL cells was indicated by the color change (D) and quantified in (E). Differentiation was induced with 2% DMSO. Both FeCl_3_ and FeEnt caused an increase in hemoglobinization as indicated by a significant change in color intensity (compared to control DMSO-treated cells). Error bars represent standard deviation. Individual data points represent technical replicates. ∗*p* < 0.05, ∗∗*p* < 0.01, ∗∗∗*p* < 0.001, ns: not significant; unpaired two-tailed Student’s *t* test.

### Ferric Ent increases differentiation and hemoglobinization of murine erythroid progenitor cells

Murine erythroleukemia (MEL) cells are murine erythroid progenitor cells that are arrested at the proerythroblast stage. In the laboratory, these cells are induced to undergo erythroid differentiation to red blood cells by addition of various chemicals (i.e. 2% DMSO) (29). Erythroid differentiation is an iron-dependent process and these cells obtain iron primarily through the transferrin transport system (DMT1 independent) (30). We tested if addition of FeEnt would positively impact the differentiation of MEL cells into red blood cells. We found that addition of FeEnt resulted in a significant increase in MEL differentiation indicated by color change (Fig. 4D-E). This increase was significantly better than supplementation with equimolar FeCl_3_ alone. Supplementing with equimolar free Ent had the opposite effect and showed a decrease in differentiation. These data suggest that FeEnt can be taken up by murine erythroid progenitor cells to a greater extent than equimolar FeCl_3_ alone, and that iron transported through this mechanism is bioavailable to promote iron-dependent differentiation.

## Discussion

Ent-mediated iron traffic was well known for its negative impact on the iron-related physiology of host animals. In animal intestinal cells, Ent produced by commensal *E. coli* was assumed to serve the bacteria in the competition for iron. Ent secreted by infectious bacteria is known to "steal" iron from the host for use by the bacteria invading the host cells (31–33). In response to infection by Ent-producing bacteria, host immune cells secrete Lipocalin2 that sequesters Ent to prevent bacterial iron scavenging (34–37). With such a reputation as a toxic factor, a beneficial role of Ent for the host seemed unlikely. A direct beneficial role of Ent on iron uptake into animal cells was first reported by a previous publication from our group (12). The finding that Ent can transport iron into the mitochondria of animal intestinal cells represents a new paradigm that points to the capability of Ent to enter mitochondria of animal cells and a new functional relationship between this siderophore molecule and animal physiology (10). A thorough understanding of the mechanism underlying the transport of FeEnt into animal cells and their mitochondria is important for us to solidify this paradigm.

In this study, we performed a set of tests in *C. elegans* and cultured mammalian cells to demonstrate that FeEnt enters animal intestinal cells through a mechanism that is independent of the well-known iron transporter DMT1. The fact that DMT1/SMF-3 is known as an apical intestinal transporter of Fe^2+^ (not Fe^3+^) may seem to eliminate it from consideration for the FeEnt mechanism, but there are three important points to consider. First, it is known that dietary ferric iron can be reduced to ferrous iron in the gut lumen by ferric reductases and subsequently transported into the cell by DMT1. Since we do not yet fully know how FeEnt is metabolized in the intestine, this was a possible scenario that needed to be tested. Second, DMT1 is also known to localize to lysosomes. While ferric iron may be transported into the cell by other means (transferrin, etc), it is subsequently delivered to the lysosome where it is then reduced to Fe^2+^. The release of this ferrous iron pool from the lysosome is partially dependent on DMT1. Loss of DMT1 would thus disrupt this step downstream of apical iron uptake. Third, DMT1 has also been found on the Mt membrane and shown to transport iron into mitochondria (16, 17). Taken together, the DMT1/SMF-3 loss of function model is deficient in all of these roles. Although DMT1 is a Fe^2+^ transporter, the lysosomal path of Fe^3+^ metabolism is intertwined with this one. It could be said that disrupting DMT1/SMF-3 function is similar to disrupting the iron endolysosomal pathway. In light of the multi-faceted functions of DMT1 in iron homeostasis, the finding that FeEnt benefits the host by a DMT1-independent mechanism is significant.

Is there another transporter that may carry FeEnt into cells? Our previous work indicated that the binding between Ent and the alpha subunit of ATP synthase (ATPSα) is required for observing the benefit of Ent-mediated iron transport. However, an *in vitro* mitochondrial iron uptake assay suggested that ATPSα, which is transported from the cytoplasm into mitochondria, is unlikely involved in the iron transport per se (12). In the search for a FeEnt transporter, a protein BLAST query of proteins similar to FepA from *E. coli* revealed that *C. elegans* lack any strong homologs. Another hypothetical mechanism would be that FeEnt may penetrate through the plasma and mitochondrial membranes by passive diffusion, which would be inconsistent with views by some siderophore experts based on what is known about the molecular weight, negative charge, and other properties of the ferric enterobactin complex (7–9). However, the diffusion model needs to be tested as it seems to be more consistent with observations made in studies by our lab and others (12, 38). It is intriguing to also consider that our data shows that this beneficial mechanism operates independently of HIF-1, ferritin, and ferroportin.

The success of Ent-mediated iron uptake in DMT-1-defective cells may suggest a potential usage of the molecule to facilitate the treatment of iron deficiency anemia (IDA), which is the most prevalent nutrient-deficiency disorder(39). Inefficient iron absorption by enterocytes can be causal to IDA and is linked to adverse side effects associated with the treatment of IDA using oral iron supplementation (39). High iron load is known to induce the expression of hepcidin, a liver-produced hormone, that binds to ferroportin to block iron efflux, thereby significantly reducing the efficacy of oral iron supplementation(39). Ent-mediated iron transport in enterocytes has the potential to elude at least some of the side effects, as the unique structure and high binding affinity of Ent-Fe^3+^ may shield Fe from interaction with certain factors that may induce undesired effects. Such a potential may be enhanced by the finding in this study that Ent-Fe^3+^ enters enterocytes and their mitochondria by a mechanism that is independent of the known import path of orally supplemented ferrous iron. Further study is certainly required to test the outstanding mechanism by which this microbial metabolite benefits the host.

## Experimental Procedures

### *C. elegans*, bacteria, and chemicals

*C. elegans* mutant strains *smf-3(ok1035)*, *smf-2(gk133), smf-1(ok1748)*, *ftn-1(ok3625), ftn- 2(ok404), fpn-1.2(ok746),* and *hif-1(ia4)* were obtained from the CGC. The Keio collection and BW25113 parental control strain were purchased from Horizon Discovery/Dharmacon. 2,2’- bipyridyl (BP) was purchased from Sigma (D216305) and dissolved in DMSO. Due to supply chain issues throughout this study, Enterobactin was purchased from Sigma (E3910), Ambeed (A570339), or purified from *E. coli* in our lab [method modified from two previous publications (40, 41)]. ^55^Fe was purchased from the National Isotope Development Center (NIDC). Bacterial cultures were grown in Luria broth-Lennox medium (10 g/L NaCl, 5 g/L yeast extract, 5 g/L tryptone, and 15 g/L agar when needed). When required, the media were supplemented with kanamycin (50 µg/mL), ampicillin (100 µg/mL), or tetracycline (50 µg/mL). Liquid bacterial cultures were grown at 37°C at 200 rpm.

### *smf-3*(-) OP50/BP growth assay

Nematode growth media (NGM) chelator plates were prepared in small batches using a scaled down version of the standard NGM recipe. Briefly, 600mL of water was combined with 1.8g NaCl, 1.5g bacto peptone, and 10.2g agar, and sterilized for 30min by autoclave. The mixture was cooled on the bench to ∼55C, then 600uL of 1M CaCl_2_, 600uL of 1M MgSO_4_, 15mL of 1M potassium phosphate buffer, and 300uL of 10mg/mL cholesterol in ethanol were added. For the BP chelator assay, 90uL of 100mM BP solution (dissolved in DMSO) was added to the agar (15uM BP final concentration). 10mL plates were poured. A fresh overnight culture of OP50 *E. coli* was used to spot the chelator plates. NGM chelator plates were stored at 4°C and used within one week for growth assays. After bacterial lawns dried, plates were supplemented with 2.5uL of 14.9mM Ent, FeCl_3_ or DMSO solvent (Fig. 1A). FeEnt was produced by incubating 14.9mM Ent with 14.9mM FeCl_3_, then adding 5uL of this FeEnt complex to each plate. 3-5 gravid adults were spot bleached on each assay plate. Worms were grown at 20°C and scored 4-5 days later, as indicated. The very same protocol was used to assay the *smf-3*(-) *ftn-1*(-)*, smf-3*(-) *ftn-2*(-), and *smf-3*(-) *fpn-1.2*(-) double mutants, and the *hif-1(ia4)* single mutant. Wild-type worms show no growth delay under these OP50/BP assay conditions (21).

### EntF(-); FepA(-) double mutant strain construction

EntF(-); FepA(-) double mutant *E. coli* strain was constructed with the lambda red recombinase system protocol (42) with minor modifications. Briefly, the *ΔEntF::Kan E. coli* BW25113 stain, which carries a kanamycin-resistant gene cassette insertion replacing the EntF gene, was selected from the Keio mutant collection (43). Its identity was confirmed by PCR genotyping with primers (CGGTGCCCTGAATGAACTGC, targeting the kanamycin-resistant gene cassette and CTGGGCGACGCCAGAGGATAATC targeting the 3’ end of the EntF gene). This strain was then transformed with the red recombinase pSIM27 vector carrying the tetracycline resistance gene cassette (Tet^r^, pSC101 *repA^ts^*) (44). Then, an ampicillin resistance cassette was amplified from plasmid pSIM8 using primers carrying at the 5′ and 3’ end 30-nucleotide extensions homologous to the *fepA* gene (Forward primer ATGAACAAGAAGATTCATTCCCTGGCCTTGTTGGTCAATCTGGGGATTTACATTCAAATATGTATCCGCTC; Reverse primer TCAGAAGTGGGTGTTTACGCTCATATACCACGTACGTCCCGGCTCGTTATAGAGTTGGTAGCTCTTGATC). PCR products were purified, then 2 µL was transformed into electro- competent *ΔEntF::Kan E. coli* cells carrying the red recombinase pSIM27 vector.

Transformants were selected on ampicillin plates and verified by colony PCR using primers (Forward primer: CGACTGCCACCAGCTCTCACTTC targeting the ampicillin resistance gene and Reverse primer: AGGGCGACACGGAAATGTTG targeting the outside region of 3’ end of the *fepA* gene) to confirm the insertion of the ampicillin resistant cassette at the *fepA* locus. The positive clone was amplified and reapplied on the LB agar plate to obtain multiple single clones. These single clones were inoculated on both plates with ampicillin/kanamycin and plates with ampicillin/kanamycin/tetracycline. We kept the clone that could not grow on tetracycline, indicating the red recombinase pSIM27 vector was lost. The *EntF*(-)*; FepA(-)* double mutant strain was tested with *C. elegans* on BP plates in the same manner described for OP50 *E. coli* above.

### *smf-3* double mutants

*smf-3(ok1035)* mutants were crossed with mutants of interest by traditional methods and confirmed by PCR.

*smf-1* PCR primers: TCATTCGGCTCTAGTGAAGGTTTGAC, GCATTGTCATGGACTGCTGATGAATC.

*smf-2* PCR primers: GTCAACACTGGCTTCATCAC, TGAGTCCGACAGTAACCTGTATTTGC *smf-3* PCR primers: GAAAGCCAAGCTACAGTAACCAGTAG, TATTATCGATTCCGCCGGAGAAGATC, CGTTATGTTCCGCGGAGATATAGTG. *ftn-1* PCR primers: ATGTGTCTCAGATTTCCGCC, GAACCCTTTCGTTGCCAATA *ftn-2* PCR primers: CGATGGAAATCTTCTAATCTGCCTGC, CTCATTGAGCAGTATTGCAGCCTTAC, TTGGTGACGACTCACGTGATATGATG *fpn-1.2* PCR primers: AGCAGAGGATTGGAGATGTAGATTGC, CAACCGAATAATCGACGAAACAGTGG, GAGAAGAGACGCAGAGATCTCATCTG

### *smf-1/2/3* knockdown test

The *smf-2(-); smf-3(-)* double mutant strain was fed *smf-1* RNAi (45) from hatching, then adults were bleached and eggs placed on the OP50/BP test plate +/- supplements [dose and preparation match the *smf-3(-)* OP50/BP growth assay detailed above]. Worms grew at 20°C and developmental stage was scored 4-5 days later, as indicated.

## Microscopy

Images of *C. elegans* were captured with a Leica M165 FC scope and Leica IC80HD camera at 2.0x magnification. Scale bar equals 500 microns.

## Cell lines and growth conditions

The human Caco-2 cell line (HTB-37) was purchased from ATCC and cultured with DMEM (Gibco 11995-040) containing 10% HI FBS (Gibco 16000-036), 100 μg/mL PEN-STREP (Lonza DE17-602E), and 1% MEM NEAA (Fisher 11140-050). HEK293F (human embryonic kidney 293-fast growing variant) cells permanently transfected with mouse FLAG-tagged (C-terminal) DMT1 2/-IRE subject to tet-on regulation have been described (28) and were maintained at 37°C with 5% CO_2_ in Dulbecco’s modified Eagle’s medium containing 10% FBS. Mouse Erythroleukemia (MEL) cells (gift from J. Brumbaugh lab) were cultured in DMEM (Gibco 11995-040) containing 10% HI FBS (Gibco 16000-036), 100 μg/mL PEN-STREP (Lonza DE17-602E), and 1% MEM NEAA (Fisher 11140-050), and maintained at 37°C with 5% CO_2_.

## Assay for ^55^Fe transport in cultured mammalian cells

Caco-2 monolayers were grown by seeding Caco-2 cells at a density of 4 x 10^4^ cells/cm^2^ in 12- well plates for about 14 days, with media changed every other day. Media was aspirated, and monolayers were rinsed with PBS. Hank’s Buffered Saline Solution (HBSS) containing 500 nM ^55^FeCl_3_ and either DMSO vehicle or enterobactin (Sigma E3910) was added to the monolayers and then incubated for 15 minutes at 37°C. 20 uM Ebselen was added 2 hours before ^55^FeCl_3_ treatment when needed. HEK293F DMT1 2/-IRE cells were grown to 70-80% confluence in 6 well plates then treated with doxycycline (50nM) (when DMT1 activity was needed) or just its solvent as control, then grown fully to confluence (24 hours). For ^55^Fe uptake experiments, HBSS containing 500 nM ^55^FeCl_3_ and either DMSO vehicle or enterobactin was added and then incubated for 15 minutes at 37°C. Fe^2+^ status was achieved by having a x10 excess of FeSO_4_ relative to the ^55^Fe and a x10 excess of ascorbic acid relative to the FeSO4. To determine intracellular ^55^Fe in both above cell lines, the media was removed, and the monolayer was rinsed with PBS (x3). The cells were then lysed with 500 μL of RIPA buffer, and radioactivity was determined on a liquid scintillation counter after diluting the cell lysate in scintillation cocktail. All values were normalized to the control cell plate unless otherwise noted.

## RNA extraction and quantitative real-time reverse transcription polymerase chain reaction (qRT-PCR)

Total RNA extraction and reverse transcription polymerase chain reaction (RT-PCR) were performed as previously described (46). The purity and concentration of RNA were spectrophotometrically evaluated by NanoDrop 2000/2000c (Thermo Fisher Scientific). A total of 2ug RNA was reverse transcribed to cDNA using the SuperScript™ IV First-Strand Synthesis System (Cat# No# 18091050 from Invitrogen, Thermo Fisher Scientific), according to the manufacturer’s guidelines. Quantitative real-time PCR (qRT-PCR) for the mouse-specific DMT1_2/-IRE was carried out in 10 µL PCR reactions using Applied Biosystems 7500 Real-Time PCR Systems using the SsoAdvanced Universal SYBR Green Supermix. The primer set for mouse −IRE DMT1 (only target on mouse DMT1 not human DMT1 presented in HEK293F cells) was 5_- ACTCCACACACTGTCCACAC -3_ (forward) and 5_GGTCTGTGTTCCCATCGAGT-3_ (reverse). The primers for β-actin as the internal control were 5_-ACTCTTCCAGCCTTCCTTCC-3_ (forward) and 5_- TGATCTCCTTCTGCATCCTGTC-3_ (reverse). Each sample was repeated in triplicate, and data analysis was performed on the Ct value of each sample using the 2^−ΔΔCt^ method with β-actin as an endogenous control for data normalization.

## MEL cell differentiation assay

MEL cells were grown in suspension, and cultured with DMEM (Gibco 11995-040) containing 10% HI FBS (Gibco 16000-036), 100 μg/mL PEN-STREP (Lonza DE17-602E), and 1% MEM NEAA (Fisher 11140-050). Media was treated with 2% DMSO to induce cell differentiation. Cells were grown in 10mL of media; *C. elegans* were grown on 10mL agar plates. Cell culture plates were supplemented in the same manner as the *C. elegans* plates, adding 2.5uL of 14.9mM Ent, FeCl_3_ or DMSO solvent to each. FeEnt was produced by incubating 14.9mM Ent with 14.9mM FeCl_3_, then adding 5uL of this FeEnt complex to each plate. Cell differentiation was scored 3-4 days later by gently pelleting the cells, removing the media, imaging the resulting pellet, and quantifying the color by ImageJ normalized to DMSO/control intensity. Three technical replicates were scored for each experiment. The beneficial effect of FeEnt was observed in three biological replicates.

## Statistical Analysis

Statistical significance was determined by t-test (unpaired, two tailed) where a p-value less than 0.05 was considered significant.

## Data availability

All data described in this study are contained within the main article.

## Acknowledgements and Funding information

We thank the CGC (funded by NIH <u>P40OD010440</u>) for worm strains, M. Garrick and J. Collins for sharing the HEK293F DMT1 2/-IRE cell line, the J. Brumbaugh lab for sharing the MEL cell line, the S. Copley lab for sharing plasmids, and PBJ Sewell and our lab members for providing helpful advice and discussion. This study was supported by the US NIH grant <u>1R35GM139631</u> (M.H.).

## Author contributions

**Aileen K. Sewell**: Conceptualization, Formal analysis, Investigation, Methodology, Validation, Visualization, Writing – original draft. **Mingxue Cui**: Conceptualization, Formal analysis, Investigation, Methodology, Validation, Visualization, Writing – review & editing. **Mengnan Zhu**: Formal analysis, Investigation, Validation. **Miranda R**. **Host**: Investigation, Validation. **Min Han**: Funding acquisition, Project administration, Supervision, Writing – review & editing.

